# Prevalence of heartwater in Guadeloupe (2024): stable endemicity and evidence of spread to Les Saintes

**DOI:** 10.1101/2025.06.11.659135

**Authors:** Dufleit Victor, Guerrini Laure, Dhune Mélanie, Viry Laetitia, Belfort Armande, Jacquet-Crétides Loïc, Deddy Jimmy, Cozier Joanna, Naves Michel, Arthein Ludovic, Samut Thony, Pezeron Manuel, Rodrigues Valerie, Damien F. Meyer, Etter Eric

**Affiliations:** CIRAD, UMR ASTRE, WOAH Reference Laboratory for Heartwater, F- 97170 Petit-Bourg, Guadeloupe, France; ASTRE, CIRAD, INRAe, Université Montpellier, Montpellier, France; Université des Antilles, École doctorale n° 636 DEECA, Campus de Fouillole, Pointe-à-Pitre – Guadeloupe, France; UR ASSET, INRAE, Centre Antilles-Guyane, Domaine de Duclos, F-97170 Petit-Bourg, Guadeloupe, France; UE PTEA, INRAE, Centre Antilles Guyane, Gardel, 97160 Le Moule, Guadeloupe, France; SANIGWA, IGUAVIE, immeuble Le Métis, LD Convenance, 97122 Baie-Mahault, Guadeloupe, France; CaribVET, site de Duclos, 97170 Petit-Bourg, Guadeloupe, France

**Keywords:** *Ehrlichia ruminantium*, heartwater, Guadeloupe, *Amblyomma variegatum*, MAP1B ELISA, seroprevalence, stochastic modelling, goat surveillance, Caribbean, One Health

## Abstract

Heartwater is a tick-borne disease caused by *Ehrlichia ruminantium* and remains endemic in the Guadeloupe archipelago. While previous data from 1989 suggested a seroprevalence of 22% in cattle, no updated figures were available for livestock, and certain islands such as Les Saintes were considered disease-free.

This study aimed to update the sero-epidemiological status of heartwater in cattle and goats across Guadeloupe in 2024. Blood samples were collected from 261 cattle and 135 goats from all islands. Serological testing using the MAP1B ELISA was performed, and true seroprevalence was estimated using a simplified stochastic model that accounts for test sensitivity, specificity, and sample uncertainty. The mean cattle seroprevalence was estimated at 28% (95% CI [22–34%]), with no significant differences between Basse-Terre, Grande-Terre, and Marie-Galante. Importantly, seropositive goats were identified on all major islands, including Les Saintes. Our findings suggest that heartwater remains endemically stable in Guadeloupe since the 1980s and reveal for the first time its serological presence in Les Saintes. These results emphasize the importance of strengthening regional surveillance systems, including veterinary reporting, serological monitoring, and coordinated efforts to prevent further spread across the Caribbean.

## Introduction

Heartwater is a severe, often fatal, infectious disease affecting wild and domestic ruminants, caused by the intracellular bacterium *Ehrlichia ruminantium*, and transmitted by the *Amblyomma* tick. It is endemic in sub-Saharan Africa, the Indian Ocean and Caribbean Islands, especially the Guadeloupe archipelago and Antigua. The tropical bont tick, *Amblyomma variegatum*, the principal vector of heartwater in the Caribbean, was introduced into the region in the 19th century and rapidly established across many islands (Barre and Uilenberg, 2010; Maillard et al., 1993). The first confirmed outbreaks of heartwater in the Caribbean were reported in the 80’s (Barré et al., 1984; Birnie et al., 1985; Perreau et al., 1980; Uilenberg et al., 1984).

The presence of the vector and the pathogen in the Caribbean has raised concerns about the potential spread to other islands and the American mainland (Kasari et al., 2010). In response, eradication programs, such as Caribbean Amblyomma Program CAP (Pegram et al., 1998), were initiated in the early 2000’s. These efforts, based primarily on systematic acaricide treatment of animals, ultimately failed to eliminate *A. variegatum* from the Caribbean region due to various reasons (Pegram et al., 2007; Ahoussou et al, 2010). In 2008, studies conducted in Guadeloupe archipelago revealed a high prevalence of *E. ruminantium* in tick populations(Molia et al., 2008; Vachiéry et al., 2008). However, the last study assessing the seroprevalence of heartwater in livestock in Guadeloupe dates back to Camus (1989) and reported 22% of heartwater seropositive cattle, though without specifying the diagnostic method used (Barré, 1989). Moreover, these studies described an endemic situation of the disease in Guadeloupe. Indeed, according to Deem et al. (1996), endemic stability is maintained by the convergence of three key factors: (1) high infection prevalence in the tick vector population, (2) the presence of reservoir hosts that sustain pathogen circulation, and (3) host infections occurring when host immunity is sufficiently high.

The present study aimed to provide an updated assessment of heartwater in the Guadeloupe archipelago in 2024 by estimating the seroprevalence in cattle and goats across all major islands, including previously non-endemic areas such as Les Saintes. Particular emphasis was placed on applying a robust statistical methodology to infer population-level prevalence while accounting for diagnostic test performance and sample uncertainty. Estimation of true seroprevalence at the population level from sampled data is a central task in veterinary epidemiology and requires inference modeling.

In epidemiology, estimating the true proportion at the population level based on a single sample relies on the assumption that the sample proportion follows a distribution. The true estimate of the mean from a single sample is calculated as 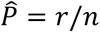, where r is the observed number of successes and n is the sample size. The standard variation of the true estimate of the mean is called the standard error, which is given by [p(1-p)/n]^1/2^. Under specific conditions, a large sample size (n>30), a prevalence within the range 0,05≤P≤0,95, and both nP and n(1-P) ≥5, the binomial distribution of the true estimate of the mean of the proportion can be approximated by a normal distribution (Thrusfield, 2007). This approach accounts for sampling uncertainty but is inaccurate for small sample size or for diseases with low prevalence. In addition, estimating the true prevalence becomes more complex when the diagnostic test used is not 100% sensitive and specific (Rogan and Gladen, 1978). Abundant literature was published to determine confidence interval that adjust for imperfect diagnostic tests (Greiner and Gardner, 2000; Lang and Reiczigel, 2014; Reiczigel et al., 2010). The adjustment of the confidence interval of the true prevalence is obtained from the adjustment of the apparent prevalence and its confidence interval, as well as from the adjustment of the sensitivity and specificity when they are estimated from samples. In this deterministic approach, the adjustments systematically use the approximation of the normal distribution. More recently, Flor et al. (2020) compared these frequentist methods with a stochastic (Bayesian) approach for estimating prevalence.

In this study, we applied a simplified stochastic approach based on Vose (2008) to estimate the distribution of the true prevalence among tested animals. This allowed to calculate the true number of infected animals in our sample. Using the same stochastic approach, we also estimated the population level seroprevalence, accounting for both the sample size and the imperfect test used, including the method used for its validation.

## Methods

### Study Area

Guadeloupe, part of the lesser Antilles and a region of the French West Indies (FWI), is an archipelago comprising five islands: Basse-Terre and Grande-Terre islands constitute the “mainland” Guadeloupe in conjunction with three smaller islands, namely La Désirade, Marie-Galante and Les Saintes. The archipelago has a predominant tropical climate, characterized by two distinct micro-climates determined by important topographic differences between the two main islands. A rainier climate is observed in Basse-Terre mostly driven by trade winds and relief (Great Soufriere volcano massif, summiting at 1 467 m) while the Grande Terre and the other islands have a dryer climate explained by absence of marked relief (Bleuse and Mandar, 1993). The livestock sector in Guadeloupe is active with around 34 800 cattle, 10 300 goat and 1 400 sheep in 2020 (Agreste, 2021). The traditional way to raise cattle in Guadeloupe is mainly tethered (Salas et al, 1986). However, this sector is declining due to various factors, including the threat caused by ticks and their associated diseases (Galan et al., 2008).

### Sampling procedure

A complete list of 4 971 breeders (34,790 head of cattle) was obtained from local authorities (DAAF Guadeloupe). Considering the intra-herd transmission due to the shared tick-infested environment, a cluster sampling of animals was decided. Based on an expected seroprevalence of 20% (Camus, 1989), and assuming a 5% error (α=0.05), an initial sample size of 246 cattle was calculated (Dohoo et al., 2014). A design effect of 1.15 was calculated based on an intra-cluster correlation coefficient of 0.05 for heartwater (Kenaw, 2023) and a cluster size corresponding to the median number of animals per breeders (n= 4). This adjustment resulted in a final required sample size of 283 cattle. To determine the numbers of farms where samples should be collected, the total number of animals was divided by the median herd size, yielding a target of 71 farms.

Sampling was conducted across the Guadeloupean archipelago from September 2023 to June 2024. The farms were sampled using a stratified sampling approach, with the proportion of farmers in each municipality serving as the sampling criterion (Dohoo et al., 2014). In some areas, convenience blood sampling was performed for goats, specifically for the purposes of disease detection. This sampling was applied in regions where cattle were absent, such as the islands of Les Saintes and La Désirade. A total of 21 goat farms with 135 animals were sampled.

Animals were sampled after obtaining farmer’s consent. The study protocol received formal ethical approval prior to implementation (APAFIS #43250-2023050210511531 v3). Blood samples were collected from restrained animals via either the jugular or the coccygeal vein, and transported under refrigerated conditions to the laboratory.

### Serological survey in cattle and goats

Serum samples from cattle and goats were tested using a modified version of the reference Elisa test for the detection of antibodies specific to the *Ehrlichia ruminantium* MAP-1B protein, a highly specific antigen previously described by Mondry et al. (1998) and (Van Vliet et al., 1995). Briefly, MAP-1B antigen was coated on microplates. After blocking, sera and positive or negative controls were incubated. Plates were washed with PBS 0,05% Tween20 and a secondary protein-HRP was added. After incubation and the last washings, TMB was added and specific colour developed if anti-MAP-1B antibodies were previously fixed. OD was measured at 450 nm and sera were declared positive if the Relative Positivity Percentage (RPP = OD sample / OD positive control x 100) exceeded 10% for goats and 24% for cattle (Rodrigues, personal communication).

### Statistical analysis

Following serological testing, cattle sera were classified as either positive or negative, indicating recent infection within the last year. Doubtful results were considered negative to prioritize specificity, as it could indicate older infections (Semu et al., 2001). Differences in seroprevalence between the three main islands, Basse-Terre, Grande-Terre and Marie Galante, were assessed using the chi-square test (χ^2^-test). Statistical significance was defined for p-values less than 0.05.

An original method incorporating uncertainty was employed to estimate the prevalence of heartwater at population level. The apparent prevalence (*AP*) was first calculated by dividing the number of seropositive animals by the total number of animals tested *(ns*). The true prevalence (*TP*) was then estimated using the standard formula (**Eq.1**), as described by Dohoo et al. (2014):

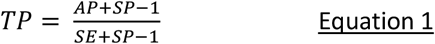

With *SP* and *SE* being the specificity and sensitivity of the MAP-1B test. To consider their uncertainty, both specificity and sensitivity were modelled using beta distributions with uniform Beta (1,1) non informative prior distribution, with parameters *shape 1* = *S + 1* and *shape 2* = *N - S + 1* (Vose, 2008). In these equations, *S* represents the number of sick animals that tested positive (for sensitivity) or the number of healthy animals that tested negative (for specificity) and *N* denotes the total number of diseased animals tested (for sensitivity) or the total number of healthy animals tested (for specificity) during the test validation study. Validation data were obtained from the WOAH reference laboratory for heartwater (Rodrigues, personal communication). The distribution of *TP* obtained was then multiplied by the number of animals tested (*ns*) resulting in a distribution of the true number of positive animals (*nsp*). To extrapolate to the entire Guadeloupean cattle population, the heartwater sero-prevalence was modelled using a beta distribution with parameter shape 1 = *nsp + 1* and shape 2 = *ns – nsp + 1*. From this distribution, the mean, 95% confidence intervals, and skewness were calculated. This method was applied to ELISA results from cattle for the whole Guadeloupe and separately for the three major islands (Basse-Terre, Grande-Terre and Marie-Galante). A classical one-dimensional Monte-Carlo simulation with 10,000 iterations was performed for stochastic modelling.

Data were stored using Excel spreadsheets (Microsoft Corporation, 2021) and statistical analysis were conducted using R software (version 4.4.3.), with the mc2d package (version 0.2.1) (Pouillot and Delignette-Muler, 2010; R Core Team, 2023).

### Geospatial interpolation

Geospatial interpolation was used to visualize the spatial distribution of heartwater across the Guadeloupe archipelago. A georeferenced layer of breeding locations was created, with each breeder represented by the centroid of their animals’ geographical positions. Farms were classified as positive if at least one animal tested seropositive for heartwater. Data from 21 goat farms were integrated into the cattle farmers’ dataset. To provide a spatial representation of heartwater seroprevalence across the Guadeloupean archipelago, Inverse Distance Weighting (IDW) interpolation was applied using a power parameter of 4 (Longley et al., 2015). This value was selected to interpolate the presence of heartwater within the Guadeloupean territory whilst ensuring the confidentiality of individual farm locations.

## Results

Sample collection was conducted on 74 cattle farms and sera were collected from 261 cattle. *Ehrlichia ruminantium* MAP-1B specific antibodies were detected in 15 out of 56 animals sampled in Basse-Terre, resulting in an apparent seroprevalence of 26.7%. In Grande-Terre, 38 out of 137 animals tested positive, yielding an apparent seroprevalence of 27.7%. In Marie-Galante, 23 of 68 animals were positives, resulting in an apparent seroprevalence of 33.8%. Overall, the combined apparent seroprevalence across all sampled cattle was 29.1% (76 out of the 261 tested cattle). Despite minor variations, the apparent seroprevalences across the three islands were comparable, and no statistically significant differences were observed between them, indicating a relatively uniform circulation of *E. ruminantium* in the cattle population across the archipelago (Table 1).

**Table 1.**
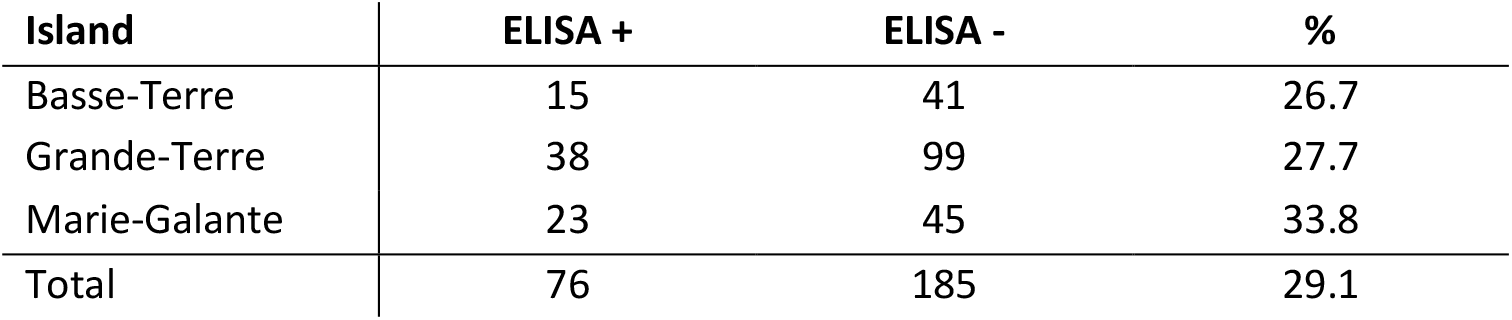
Seroprevalence of *Ehrlichia ruminantium* in cattle by island: results of the MAP1-B ELISA test performed on cattle sampled from the three major islands of the Guadeloupe archipelago. A total of 261 animals were tested: 56 from Basse-Terre, 137 from Grande-Terre, and 68 from Marie-Galante.

The distribution of cattle true seroprevalence modelled at population level presented a mean of 0.28 with a 95% confidence interval of [0.22; 0.34]_95%_ and a skewness of - 0,01 for the whole Guadeloupe, 0.27 [0.16; 0.39]_95%_ with a skewness of 0.26 for Basse-Terre, 0.27 [0.20; 0.36] _95%_ with a skewness of 0.16 for Grande-Terre and 0.34 [0.23; 0.45] _95%_ with a skewness of 0.11 for Marie-Galante. Taken together, these results support the conclusion that heartwater seroprevalence remains consistently endemic across the Guadeloupe archipelago without marked geographical heterogeneity. (**Figure 1)**.

**Figure 1.**
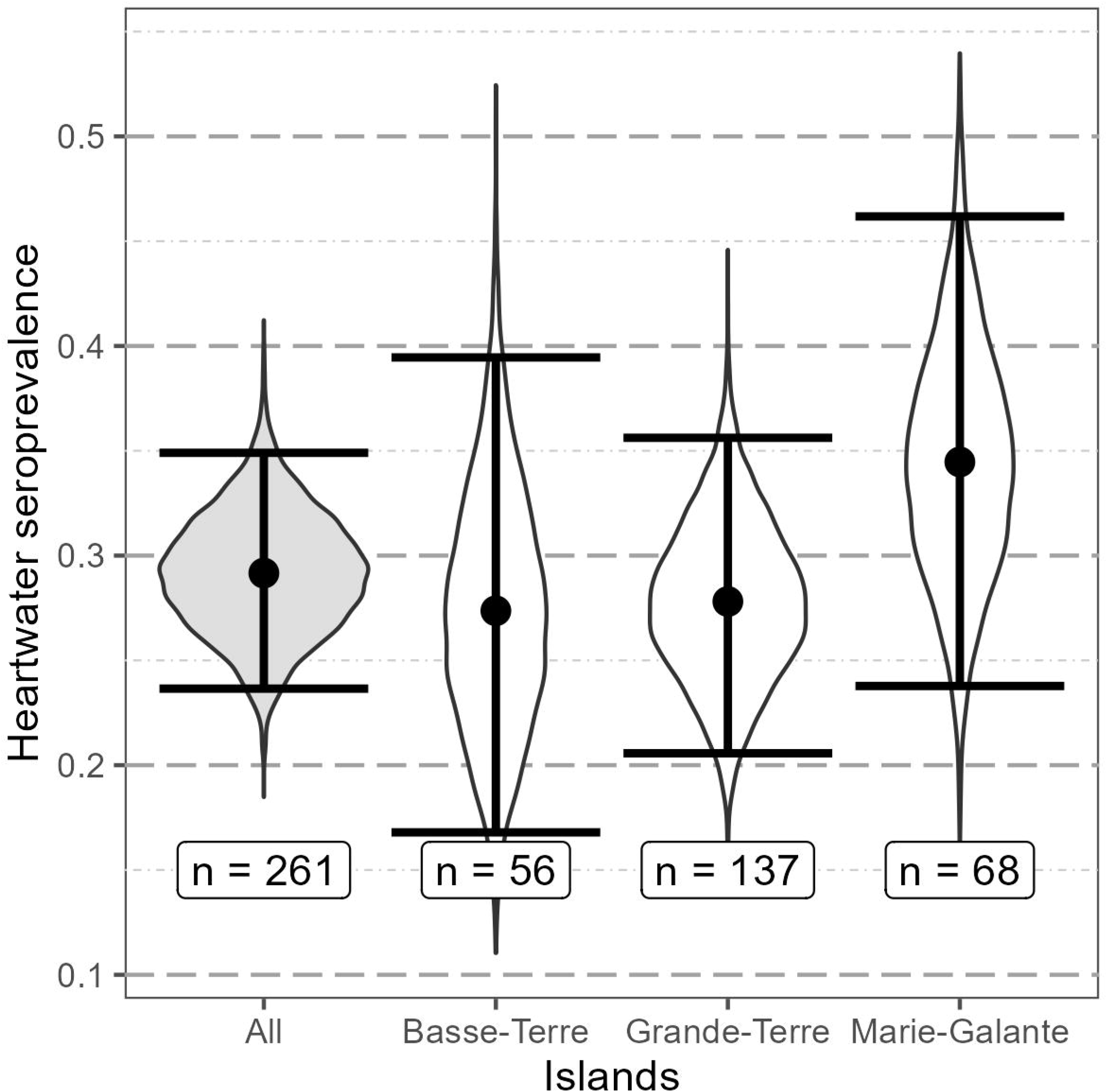
Modelled heartwater seroprevalence in cattle across the Guadeloupe archipelago. The violin plot displays the beta-distributed estimates of *Ehrlichia ruminantium* seroprevalence in cattle, modelled for the main islands of the Guadeloupe archipelago—Basse-Terre, Grande-Terre, and Marie-Galante—as well as the combined dataset (“All”). Each distribution was derived from stochastic modelling, incorporating diagnostic test uncertainty and sample size. Dots represent the mean seroprevalence, bars indicate the 95% confidence interval, sample sizes (n) are indicated for each group. The density curves are depicted by the shapes around the centre lines.

Amongst the 74 cattle farms selected, 47 had at least one animal that tested positive for *E. ruminantium* using the MAP-1B serological test. The MAP-1B ELISA was also performed on 135 goats’ serum samples collected from 21 farms located in Basse-Terre, Grande-Terre, Marie, Galante, la Désirade, and les Saintes. Amongst them, 27 seropositive animals were detected on all investigated islands, further supporting the broad geographic distribution of heartwater among ruminants in Guadeloupe.

The inverse distance weighting map further supported the widespread presence of heartwater, showing evidence of infection on farms across all islands within the archipelago (Figure 2). Seropositive farms (goats or cattle) were detected across all major islands, including Basse-Terre, Grande-Terre, Marie-Galante, La Désirade, and, notably, Les Saintes. High seropositivity zones were especially concentrated in the central and northern parts of Grande-Terre, the southwestern and eastern parts of Marie-Galante, and several lowland regions of Basse-Terre (Figure 2). Municipalities not included in the sampling are depicted in grey on the map, while the Guadeloupe National Park is highlighted in dark green. As expected, no samples were collected from within the National Park boundaries due to livestock absence in these protected areas. The spatial distribution of seropositive farms confirms the widespread circulation of *Ehrlichia ruminantium* throughout the archipelago, with no major exclusion zones identified.

**Figure 2.**
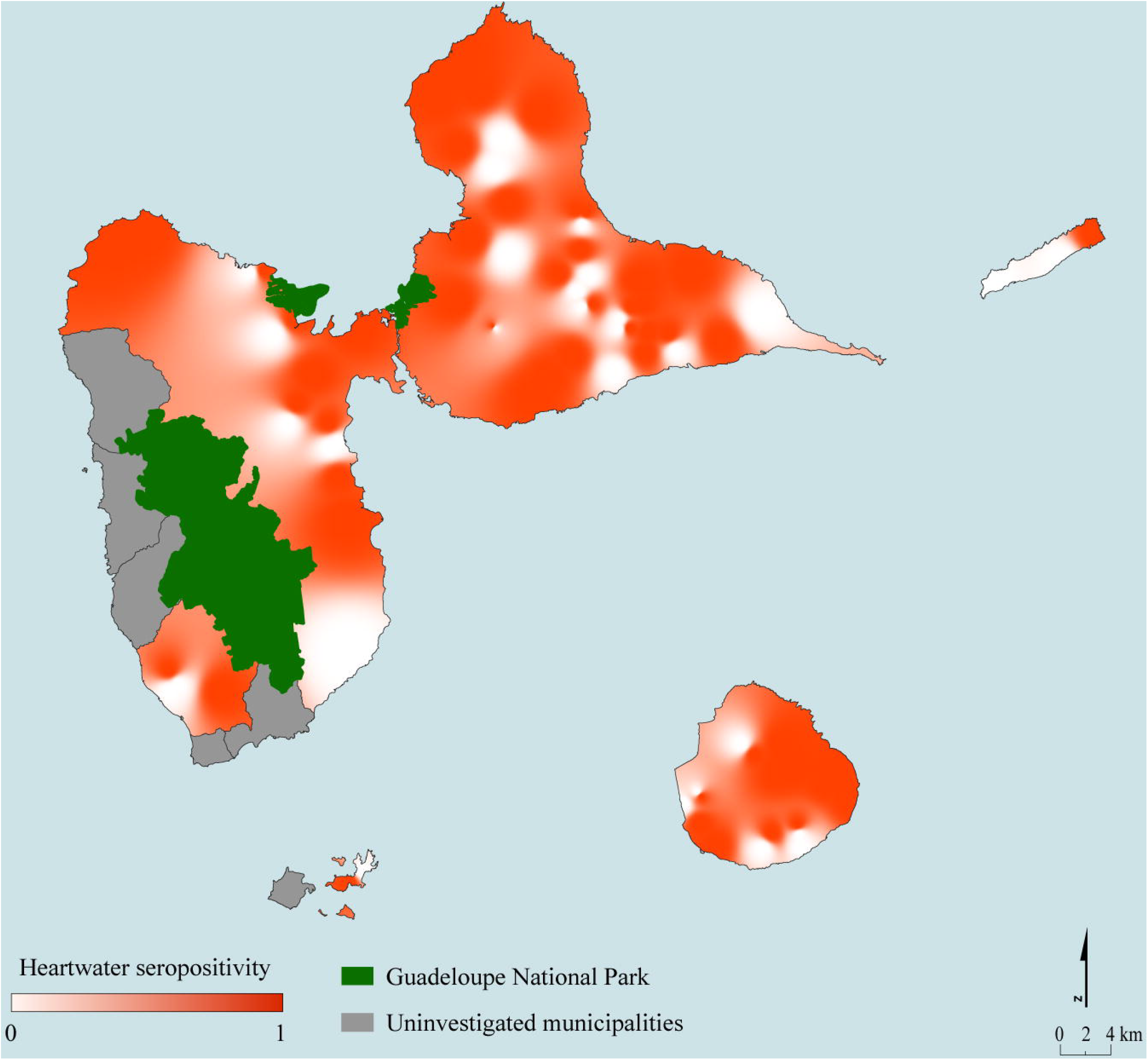
Spatial distribution of heartwater seropositivity in cattle and goats across the Guadeloupe archipelago. This map presents the interpolated spatial distribution of heartwater seropositivity based on MAP 1B ELISA results from cattle and goat farms. Inverse Distance Weighting (IDW) interpolation was applied to farm-level data to visualize the relative intensity of seropositivity across the region. The color gradient ranges from white (no seropositivity, value = 0) to deep red (seropositivity evidence, value = 1).

## Discussion

The results of this study revealed a heartwater seroprevalence of 0.28 (95% CI [0.22; 0.34]) in cattle across the Guadeloupe archipelago. The simplified stochastic method to infer population-level seroprevalence produced a relatively symmetric distribution for the whole Guadeloupe, and slightly right-skewed distributions for individual islands. However, statistical analysis of the raw data did not reveal any significant differences in apparent seroprevalence between Basse-Terre, Grande-Terre, and Marie-Galante. These results are consistent with the 22% seroprevalence reported by Camus in 1989 (lower limit of the current confidence interval) reinforcing the hypothesis of epidemiological stability of *Ehrlichia ruminantium* circulation in Guadeloupe over the past 35 years.

The consistent circulation of *Ehrlichia ruminantium* in Guadeloupean cattle suggests that heartwater is endemically stable, potentially maintained by repeated exposure. Similar results have been reported among cattle in endemic regions of West and Southern Africa, where seroprevalence detected using the MAP-1B ELISA test range from 20% to 40% (Bell-Sakyi et al., 2003; Mahan et al., 1998; Peter et al., 2001). The transient nature of the antibody response to the MAP-1B antigen, especially under conditions of recurrent natural exposures, may result in moderate but stable seroprevalence levels over time (Semu et al, 2001). Therefore, these data provide strong evidence that *E. ruminantium* was actively circulating in the Guadeloupean cattle population during 2023-2024.

Serological evidence of heartwater was also detected in goats across all surveyed islands, including La Désirade and Les Saintes. The detection of seropositive animals in Les Saintes, a region historically considered free of heartwater, suggests a recent introduction and ongoing spread of the pathogen beyond previously recognized endemic areas. Based on veterinary field investigations and animal movement records, the most plausible hypothesis is the introduction in 2015 of infected livestock or animals infested with infected ticks from Marie-Galante. End 2024, nervous clinical signs on goats were reported by farmers and the mandated veterinarian then after confirmed by molecular diagnosis (unpublished data) supporting the active presence of *E. ruminantium* in Les Saintes. These findings highlight the successful establishment and active dissemination of the disease across all major islands of Guadeloupe. While many other Caribbean islands are currently infested with *Amblyomma variegatum* ticks (Charles et al., 2020; Kasari et al., 2010), most remain free of heartwater, as evidence by the absence of reported clinical signs or molecular detection of *Ehrlichia ruminantium*. The last study on sheep, cattle, and goats from various Caribbean islands, conducted in 2011, reported less than 1% seroprevalence with no clinical cases, but these findings were then attributed to false positive reactions probably due to cross-reactivity of MAP-1B antigen with other tick-borne ehrlichiosis (Kelly et al., 2011).

The recent detection of seropositive goats in Les Saintes, an area previously considered free from heartwater, underscores the need to maintain targeted surveillance in the region. Establishing a robust syndromic surveillance system, involving veterinary practitioners, is critical for the early detection of heartwater outbreaks and for tracking seasonal or annual variations in disease prevalence (Thompson and Etter, 2015). By systematically recording clinical cases, tick infestations, suspected heartwater occurrences, seroprevalence and *Ehrlichia ruminantium* detection in ticks, veterinary services could provide valuable epidemiological data to inform both preventive and control strategies. In Guadeloupe, such a program was implemented with the RESPANG project in 2010 (Vachiéry et al., 2011). An innovative surveillance network was established to collect and diagnose ruminants showing nervous syndromes. This network involved veterinary practicians, research sector and local authorities (Lefrancois et al., 2011), but unfortunately, RESPANG is not active anymore.

The spread of heartwater in Guadeloupe archipelago highlighted the necessary cooperation between Caribbean islands. This collaboration should focus on understanding the introduction dynamics and tracking the spread of infected ticks and/or livestock between islands. Potential risk factors analysis would improve recommendations to control heartwater and mitigate tick infestations. A coordinated effort involving local farmers, veterinary services, and epidemiologists will be essential to prevent further spread of the disease to neighbouring islands. Such cooperation already existed in the Caribbean with the Caribbean Amblyomma Program (CAP) (Barré et al., 1996; Pegram et al., 1998) which aimed to eradicate *Amblyomma variegatum* from the Caribbean but failed due to numerous factors (Pegram et al., 2004, 2007; Ahoussou et al, 2010). Today, the *CaribVET* ^1^network is ideally positioned to coordinate and facilitate such initiatives. Leveraging its established connections with veterinary authorities and research institutions across the Caribbean, *CaribVET* could promote standardized protocols for sampling, diagnosis, and data sharing. This coordinated approach would strengthen the region’s ability to early detect outbreaks, assess epidemiological trends, and implement effective control measures. Despite existing vector control measures promoted by the Guadeloupean sanitary defence group (SANIGWA), the persistence of *Amblyomma variegatum* populations and the continued spread of heartwater underscore the need for more effective tick management strategies. Moreover, accelerating the development of a heartwater vaccine remains a key priority if long-term control of this economically significant disease is to be achieved.

## Conclusion

This study confirmed that heartwater remains endemic in Guadeloupe. Those results represent the first estimation of Guadeloupean cattle seroprevalence using the WOAH reference ELISA. An original statistical method was applied to determine heartwater prevalence at the cattle population level. Stable levels of seroprevalence have been observed on the main islands. Furthermore, there is evidence to suggest that the virus has spread to previously unaffected areas, such as Les Saintes. These findings underscore the need for strengthened regional surveillance, control strategies and partnership between actors of the livestock sector. Reactivation of local network, such as the previously existing RESPANG, appear determinants to efficiently collect valuable data on disease’s epidemiology.

Given the continued presence of *Amblyomma variegatum* across the Caribbean and the risk of further geographic expansion, coordinated efforts in tick management and disease prevention at a regional scale are essential. Investment in research, including vaccine development and regional monitoring, will be critical to safeguarding livestock health and maintaining agricultural resilience across the Caribbean and beyond. By combining local and regional surveillance, stakeholders can better anticipate epidemiological trends, limit disease spread and safeguard livestock industries throughout the Caribbean region.

## Conflict of interest disclosure

The authors declare that they have no financial conflicts of interest in relation to the content of the article.

## Funding

This work is supported by the United States Department of Agriculture grant 58-3022-1-018-F (Risk of Arthropod-borne diseases in the Caribbean). The authors also thank the Guadeloupe region and European Agricultural Fund for Rural Development (TISARU project, FEADER_M16_2021_01) for providing logistical support in the sampling of animals.

## Acknowledgements

Firstly, we would like to express our sincere gratitude to all the farmers who participated in our survey. Thanks to Dr Gil MANUEL, who’s first reported potential heartwater cases in Les Saintes and for the facilitation of our sampling in this area. We also thanks Sylvie CHAUMIEN-LECOLLINET, Jean Christophe BAMBOU, Alain FARANT, Xavier GODARD, Tony KANDASSAMY, Ferdy NIMIRF, Frédéric POMMIER, Nathan RANGASSAMY, Dimitri ROMIL-GRANVILL for the help provided during specific caprine sampling.

## Author contributions

Dufleit Victor: Data curation, Formal analysis, Investigation, Methodology, Software, Validation, Visualization, Writing – original draft ; Guerrini Laure: Formal analysis, Methodology, Visualization, writing – review and editing ; Dhune Mélanie: Investigation, Methodology, Resources ; Viry Laetitia: Investigation, Methodology, Resources ; Belfort Armande: Investigation, Methodology, Resources ; Jacquet-Crétides Loïc: Investigation ; Deddy Jimmy: Investigation ; Cozier Joanna: Investigation, Data curation, Writing – review and editing ; Naves Michel: Conceptualization, Investigation, Methodology, Funding acquisition, Supervision, Writing – review and editing ; Arthein Ludovic: Investigation ; Samut Thony: Conceptualization, Investigation, Funding acquisition ; Pezeron Manuel: Conceptualization, Investigation, Funding acquisition ; Rodrigues Valerie: Conceptualization, Methodology, Writing – original draft, Supervision, Writing – review and editing ; Meyer Damien F.: Conceptualization, Methodology, Supervision, Writing – original draft, Writing – review and editing ; Etter Eric: Conceptualization, Methodology, Validation, Supervision, Writing – original draft, Writing – review and editing.

## Footnotes

1 https://www.caribvet.net/

## Notes

### Competing Interest Statement

The authors have declared no competing interest.

### Summary of Updates

We have submited a revision because we forgot to mention an important person in "Aknowledgment".

